# Genetic architecture and selective sweeps after polygenic adaptation to distant trait optima

**DOI:** 10.1101/313247

**Authors:** Markus G Stetter, Kevin Thornton, Jeffrey Ross-Ibarra

## Abstract

Understanding the genetic basis of phenotypic adaptation to changing environments is an essential goal of population and quantitative genetics. While technological advances now allow interrogation of genome-wide genotyping data in large panels, our understanding of the process of polygenic adaptation is still limited. To address this limitation, we use extensive forward-time simulation to explore the impacts of variation in demography, trait genetics, and selection on the rate and mode of adaptation and the resulting genetic architecture. We simulate a population adapting to an optimum shift, modeling sequence variation for 20 QTL for each of 12 different demographies for 100 different traits varying in the effect size distribution of new mutations, the strength of stabilizing selection, and the contribution of the genomic background. We then use random forest regression approaches to learn the relative importance of input parameters in determining a number of aspects of the process of adaptation including the speed of adaptation, the relative frequency of hard sweeps and sweeps from standing variation, or the final genetic architecture of the trait. We find that selective sweeps occur even for traits under relatively weak selection and where the genetic background explains most of the variation. Though most sweeps occur from variation segregating in the ancestral population, new mutations can be important for traits under strong stabilizing selection that undergo a large optimum shift. We also show that population bottlenecks and expansion impact overall genetic variation as well as the relative importance of sweeps from standing variation and the speed with which adaptation can occur. We then compare our results to two traits under selection during maize domestication, showing that our simulations qualitatively recapitulate differences between them. Overall, our results underscore the complex population genetics of individual loci in even relatively simple quantitative trait models, but provide a glimpse into the factors that drive this complexity and the potential of these approaches for understanding polygenic adaptation.

**Author summary:** Many traits are controlled by a large number of genes, and environmental changes can lead to shifts in trait optima. How populations adapt to these shifts depends on a number of parameters including the genetic basis of the trait as well as population demography. We simulate a number of trait architectures and population histories to study the genetics of adaptation to distant trait optima. We find that selective sweeps occur even in traits under relatively weak selection and our machine learning analyses find that demography and the effect sizes of mutations have the largest influence on genetic variation after adaptation. Maize domestication is a well suited model for trait adaptation accompanied by demographic changes. We show how two example traits under a maize specific demography adapt to a distant optimum and demonstrate that polygenic adaptation is a well suited model for crop domestication even for traits with major effect loci.

## Introduction

### Adaptation

Understanding molecular adaptation is essential for the study of evolutionary processes, genetic diseases, and plant and animal breeding. The process of adaptation is often divided into three separate modes: hard selective sweeps, soft selective sweeps and polygenic adaptation [1]. In recent decades many empirical population genetic analysis have focused on hard selective sweeps because these leave a distinct molecular signature that can be readily detected in genomic data. Hard sweeps result from the reduction of genetic diversity at neutral sites linked to a new beneficial mutation that rapidly fixes [2]. In recent years, other forms of selection that play an important role in evolution and adaptation have begun to receive increased attention, although these are more difficult to detect in empirical data. For instance, sweeps from selection on standing genetic variation leave a less distinct pattern on diversity than hard selective sweeps because the beneficial variant has had more time to recombine onto multiple genetic backgrounds [3, 4]. In addition to processes involving sweeps at individual loci, polygenic adaptation — in which selection acts on a quantitative trait with complex genetic architecture — is frequently regarded as a third mode of adaptation and can lead to rapid phenotypic change via relatively minor shifts in allele frequencies [5].

Although well-studied traits such as human height [6], coat color in mice [7] and grain yield in crops [8] follow patterns consistent with the polygenic pattern, the dynamics and genetic architecture of polygenic adaptation are not well understood. Polygenic adaptation has only gained importance in empirical population genetics relatively recently, but the field of quantitative genetics is based on the idea that traits are controlled by large numbers of loci [9]. Population genetics and quantitative genetics diverged with the appearance of the first molecular data allowing empirical evaluation of single locus population genetic models, while the analysis of effects of single loci in quantitative genetics has long been limited by the large number of phenotyped individuals needed [10]. The increasing availability of high density SNP sets and whole genome sequencing for tens of thousands of individuals, however, is now providing the opportunity to test both population and quantitative evolutionary genetic hypotheses in empirical data [e.g. 11].

Many polygenic traits are thought to evolve under stabilizing selection, in which selection acts against extreme deviations from an optimum trait value [12, 13, 14]. Under such a model, an individual’s fitness is given by its phenotypic distance from the trait optimum and the strength of stabilizing selection. Within this framework, recent attention has focused on the dynamics of polygenic adaptation to a new nearby phenotypic optimum [15, 16, 17, 18, 19, 9]. In this scenario, genetic variance in the population decreases when most effect sizes are small, because many sites fix. In contrast, when most mutations have large effect sizes, the genetic variance increases because large effect loci increase in frequency but do not fix [15, 19]. In addition to allele frequency changes, theoretical quantitative genetic analyses have revealed that selective sweeps are prevalent during polygenic adaptation [20, 17]. These studies have developed important theoretical background for the understanding of polygenic adaptation and have documented the dynamics of a small number of loci during the course of adaptation. Each of these studies shows in detail how a small number of parameters influences adaptation, but the complex interplay of mutation, selection, and demography across a large parameter space has not yet been explored. For example, population growth has been shown to influence the contribution of low frequency alleles to trait variance [21], but the interaction of demography with parameters such as the distribution of effect sizes of new mutations needs further investigation.

Here, we take a simulation approach to study a population adapting to an optimum shift, modeling sequence variation for 20 QTL for each of 12 different demographic models for 100 different traits with varying effect size distribution of new mutations, strength of stabilizing selection, and the contribution of the genomic background beyond the simulated QTL. After detailed analysis of a single scenario, we use machine learning to extract parameter importance for the input parameters. Our results illustrate that selective sweeps are common under most scenarios, even for mutations of relatively minor effect. We employ machine learning on genetic architecture matrices and find that demography and the effect size of new mutations have the largest influence on present day genetic architecture. After identifying general parameter importance, we use maize domestication as an example and investigate two diverging traits in a population that underwent a population bottleneck and exponential growth [22], showing how these traits adapt to the changing optimum and comparing our findings to archaeological and genetic data [23, 24].

## Results

We first simulated adaptive and stabilizing selection on a single quantitative trait in a randomly mating diploid population. After a burn-in to equilibrium, we simulated an instantaneous shift in the optimal trait value from 0 to 10, corresponding to 89.6 z-scores of 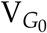. The population underwent truncation selection until reaching the new optimum, at which time stabilizing selection resumed. We assumed an additive model with no epistasis, and simulated 20 unlinked QTL as well as a genomic “background” over a range of parameters describing population demography and the trait, including the effect size of new mutations, strength of stabilizing selection, distance to the new optimum, effects of genomic background and population, and bottleneck severity and population expansion (Table 1).

**Table 1.** Parameters and variables.

| Variable | Description |
| --- | --- |
| $N_{\text{anc}}$ | Population size at equilibrium |
| $N_{\text{final}}$ | Population size after $0.1 \times N_{\text{anc}}$ generations |
| $N_{\text{bottleneck}}$ | Population size during bottleneck |
| $\psi$ | Proportion of phenotype due to genetic background outside of QTL |
| $\sigma_m$ | Standard deviation of effect sizes of new mutations |
| $V_S$ | Strength of stabilizing selection |
| $V_{G_0}$ | Genetic variance at equilibrium |

### Single simulation results

The adaptation of a quantitative trait to a sudden environmental change involves allele frequency shifts at many sites, some of which result in selective sweeps. To build intuition around basic patterns seen in these simulations as a population adapts to a new optimum, we first describe results of a single simulation with constant population size, intermediate effect sizes of new mutations (*σ*_*m*_=0.05), strong stabilizing selection (*V*_*S*_=1), and no phenotypic effect of the genomic background (*ψ*=0). We present how such a population adapts to the new optimum and how allele frequencies and effect sizes change during this process (Figure 1).

**Figure 1.**
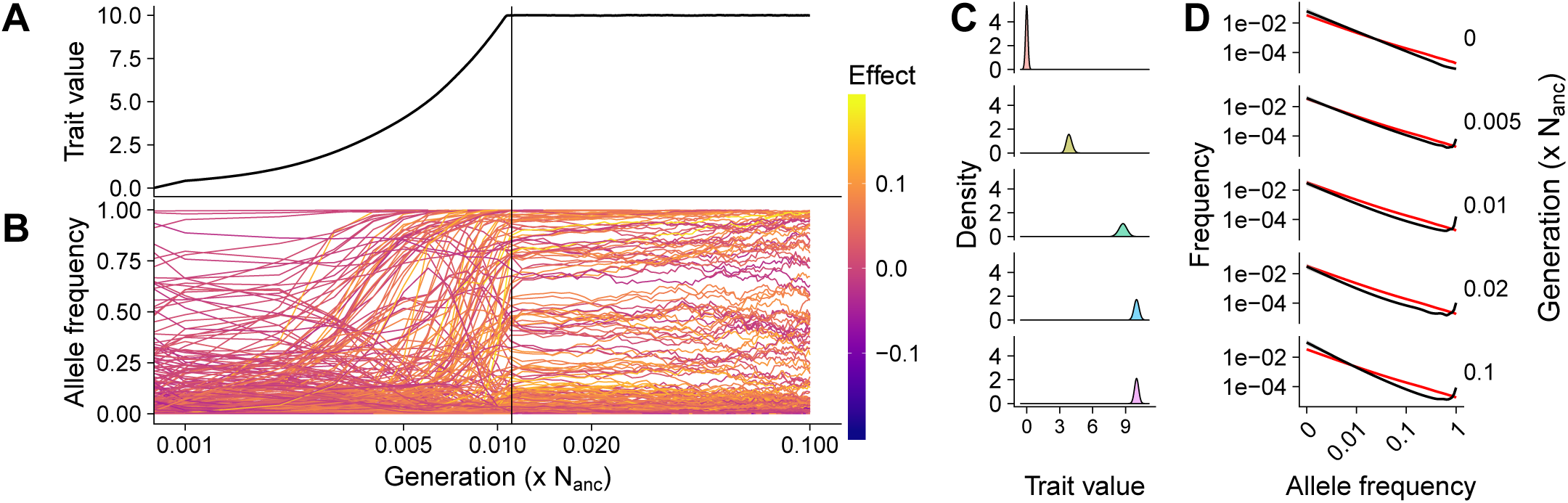
Population dynamics of a single parameter set. **A**) Trait evolution after an optimum shift and **B**) allele frequency dynamics during adaptation from a single replicate. The vertical line shows when the new trait optimum was reached and line colors denote effect sizes, and time is shown on a log scale. **C**) The phenotypic distribution and **D**) site frequency spectra of segregating mutations (black) and neutral expectation (red) from 100 independent replicates. Panels show different generations including equilibrium prior to adaptation (0), during adaptation (0.005), just before the new optimum is reached (0.01), after the new optimum has been reached (0.2), and the final generation (0.1). All results are from a simulated population with constant population size, *σ*_*m*_ = 0.05, and *V*_*S*_ = 1.

The population mean trait value increased linearly (*log*_10_ scaled x-axis in figure 1A) until shortly before the new optimum was reached within 0.011 (sd= 0.0004) N_anc_ generations (Figure 1A and C). As the population mean approached the optimum the rate of change decelerated, presumably because some individuals now had phenotype values above the optimum such that alleles which contribute positively to the trait are no longer uniformly beneficial to fitness. The trait variance increased after the optimum shift and during the adaptation process. Though it declined once the new optimum was reached, it did not return to the equilibrium variance by the end of the simulation (Figure 1C). This increase in variance is generated by the increase in allele frequency of formerly rare, large positive effect alleles.

Following individual mutations shows that, at the onset of the optimum shift (generation 0) alleles with negative effect sizes rapidly decline in frequency unless they were already near fixation (Figure 1B). Alleles with positive effects, on the other hand, increase quickly in frequency and fix. Once the new optimum is reached, frequencies of both positive and negative alleles changed slowly, but the number of small effect alleles increased. This shows how a population can adapt to a sudden environmental change by an increase of beneficial alleles and decrease of disadvantageous alleles in a relatively short time.

Looking at the change in allele frequencies of all mutations helps to understand what drove the adaptation process in the population (Figure 1D). At equilibrium, variants with larger effects are selected against, leading to an excess of rare variants compared to neutral expectations. The site frequency spectrum (SFS) then changed quickly after the optimum shift as selection fixed positive mutations. Directly before the new optimum was reached (0.01N_anc_), 11% of mutations were at very high frequencies (*>* 0.5) while after reaching the new optimum (0.02N_anc_) only 8% of mutations were at such high frequencies and the number of high frequency segregating sites further declined in consecutive generations. Under stabilizing selection, extreme values are again selected against and alleles that have risen to intermediate frequency during adaptation return to their equilibrium frequency. By 0.1 ×N_anc_ generations the SFS again reflected an excess of rare alleles, but also an excess of high frequency derived alleles. The observed high frequency derived alleles ((Figure 1D) bottom) represent in fact their ancestral counterpart, which is at low frequency. These mutations increased in frequency during adaptation, but both alleles have the same fitness effect after the equilibrium has been reached and the mutation does consequently not decrease in frequency.

When a selected mutation increases in frequency quickly, it often reduces diversity in adjacent genomic regions, leading to a pattern commonly referred to as a selective sweep. While we cannot assess diversity at linked neutral sites in our model, we can nonetheless identify likely selective sweeps by comparing the sojourn time of individual alleles to that of a neutral allele experiencing equivalent demographic processes (see Methods). Following these criteria, 72% of all fixations in this simulation were selective sweeps. Of these, 73% were sweeps from standing variation. While there was an overall negative correlation between the time a site was segregating in the population and its effect size on the trait, there were a number of mutations that fixed later than expected given their effect size (Figure 2A).

**Figure 2.**
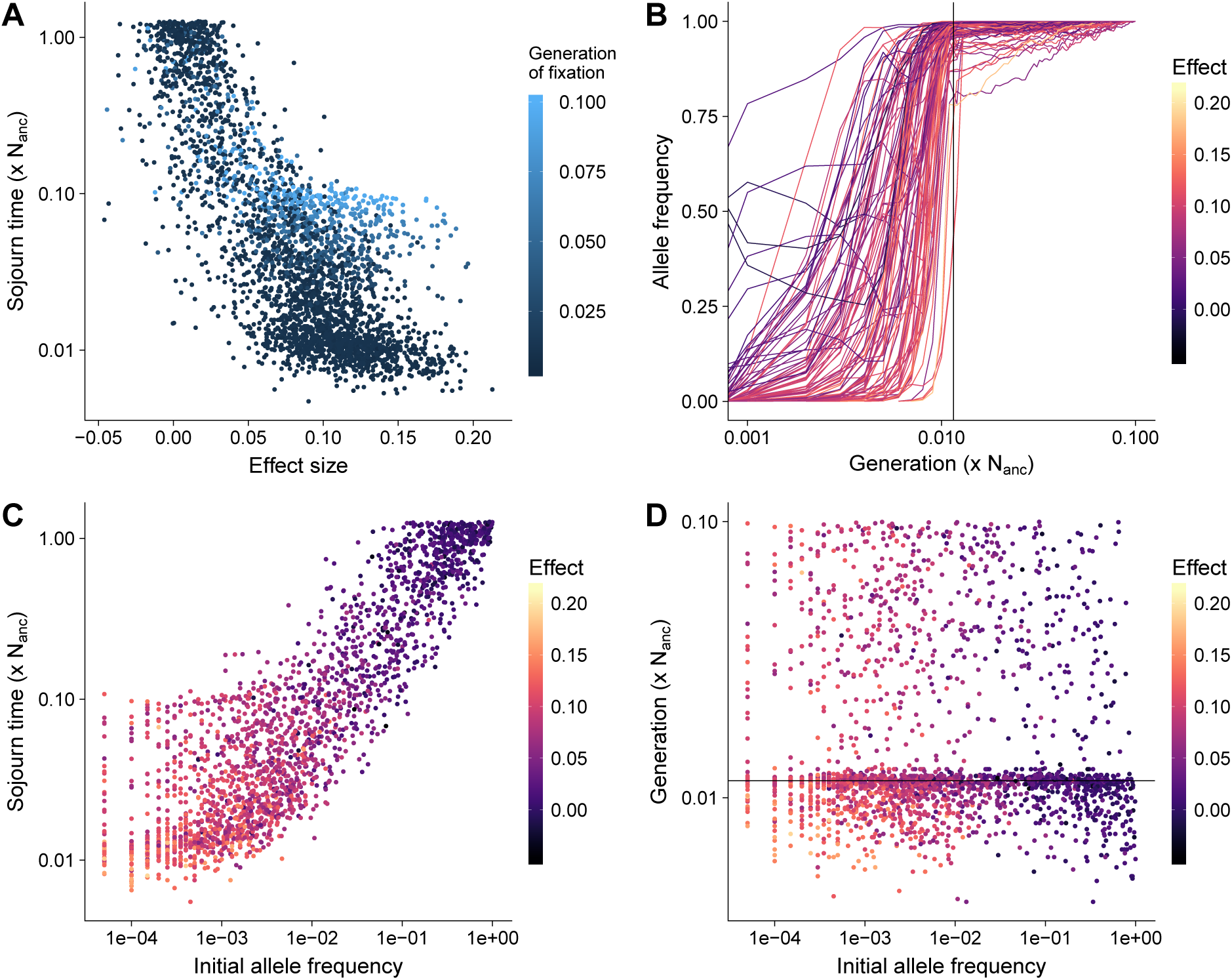
Selective sweeps. **A**) Speed of fixation of selective sweep mutations. **B**) Dynamics of fixations that occur after the new optimum was reached. **C**) Speed of fixation of sweeps from standing variation compared to their initial frequency. **D**) The generation at which sweeps from standing variation fix. All results are from a simulated population with constant population size, *σ*_*m*_ = 0.05, and *V*_*S*_ = 1, and time is shown on a log scale.

Observing the frequency trajectories of sites that fixed after the new optimum had been reached shows that the speed of frequency change for sites that fix after the new optimum had been reached slowed down substantially, but they eventually reached fixation. When the new optimum has been reached, any increase or decrease in frequency of large effect mutations takes the population away from the trait optimum and is selected against. The remaining change in frequency is mostly stochastic and results from minor fluctuations in the trait mean due to frequency changes at other sites [19]. However, because stabilizing selection acts against stochastic variation in allele frequencies that move the population away from the optimum, the time to fixation or loss for an allele is still faster than neutrality in a manner that has sometimes been deemed similar to underdominance [25]. Some mutations with negative effects that decreased in frequency under truncation selection after the optimum shift can then increase in frequency again once the new optimum is reached and stabilizing selection takes over (Figure 2B). Such mutations provide a good example of selection on a quantitative trait, which results in selection coefficients that can vary in sign or magnitude depending on the total phenotypic value of the individual in which they occur, its distance to the optimum, and the details of when and what kind of selection occurred.

In our simulations, fixations from standing variation fixed either fast, because they were present at high frequency at the onset of directional selection, or due to their large effect on the trait. However, there was no correlation between the initial allele frequency and the generation in which the mutation fixed (Figure 2C and D). Large effect mutations segregated at low frequency in the equilibrium population, while small effect sites were already at higher frequencies, explaining why large effect and small effect mutations fixed at similar generations, despite the difference in speed of allele frequency shift. Negative and effectively neutral mutations may also fix together with large effect positive mutations presumably due to the effects of genetic hitchhiking (Figure 2).

### Complex genetic architectures with demographic changes

The detailed analysis of a single population adapting to a sudden environmental change helps to build intuition on the dynamics of a specific set parameters, but is far from the complexities of quantitative trait evolution in natural populations. For example, most populations have experienced some form of fluctuation in population size, and traits differ both in the strength of stabilizing selection as well as in their genetic architecture — the frequency and effect size of mutations that cause variation in the phenotype. To understand the effect of these and other variables, we simulated 1,200 different combinations of parameter sets to examine the contribution of the strength of stabilizing selection, the effect size of new mutations, population demography, and differences in genetic background on variation and adaptation of the focal trait.

The combination of *V*_*S*_ and *σ*_*m*_ led to different genetic variances at equilibrium ranging from 0.004 to 0.751, leading to a distance of the new trait optimum between 11.5 and 158.2 z-scores (Figure S3). We calculate *V*_*G*_ in every generation during the burn-in and compared it to the expected genetic variance approximated under the House of Cards (HoC) or stochastic HoC [26, 27] approximations [12]. The majority of simulations are within the regime of HoC, though the approximation underestimated *V*_*G*_ for *σ*_*m*_ = 0.9 and *V*_*S*_ = 1 and overestimated *V*_*G*_ for large *V*_*S*_ and small *σ*_*m*_; all simulations closely expetations under the stochastic HoC approximation. All burn-ins had a mean fitness close to one at equilibrium after 10*N* generations and the mean *V*_*G*_ was constant (Figure S4 and S3).

To understand the factors driving variation in particular aspects of the data, we employed a random forest machine learning model (see Methods) to retrieve parameter importance.

### Speed of polygenic adaptation

An important factor for the survival of a population exposed to changing environments is how fast it can adapt to new conditions. Our simulated populations varied widely in the time required to reach the new optimum, from 0.001 to 0.99 N_anc_ generations. A total of 732 of the 120,000 simulations did not reach the new trait optimum within the simulated time of 0.1 ×N_anc_ generations, but all parameter combinations had at least 8 (of 100) replicates reaching the new optimum. In general, simulations that did not reach the new optimum were those with a strong bottleneck (reduction to 1% or 5% of N_anc_). In particular, more than 70% of all simulations with the smallest *σ*_*m*_ (0.01), no genetic background, 1% bottleneck, and a final size of N_anc_ did not reach the new optimum, regardless of their strength of stabilizing selection (*V*_*S*_).

All three adaptation-related summary statistics (time to new optimum, adaptation rate, and *V*_*G*_ in the final generation) were well predicted, with cross-validation accuracy over 90%. Overall, the parameter contributing most to this variation is *σ*_*m*_, with a relative importance of *>* 50% (Figure 4). This was followed closely by the proportion of the trait explained by genetic background (*ψ*) at 31%, while demography and *V*_*S*_ were of relatively minor importance (Figure 3 and S5). We find that the rate of phenotypic adaptation was highest for populations with small *σ*_*m*_ and large *ψ*, and these two factors explained the majority of the observed variation (Figure 4). The initial genetic variance, a combination of *V*_*S*_ and *σ*_*m*_, was the best predictor for the genetic variance in the final generation, but the strength of the bottleneck and *ψ* had a relative importance of 11% and 17%, respectively (Figure S5). The genetic variance in the final generation increased with increasing *σ*_*m*_, though it plateaued at the largest *σ*_*m*_ simulated (Figure 4). Of the 1,200 parameter combinations, 45 had a mean *V*_*G*_ as high as the initial *V*_*G*_ (*±* 5%), 410 had higher *V*_*G*_ and 735 had lower *V*_*G*_ than the initial equilibrium population.

**Figure 3.**
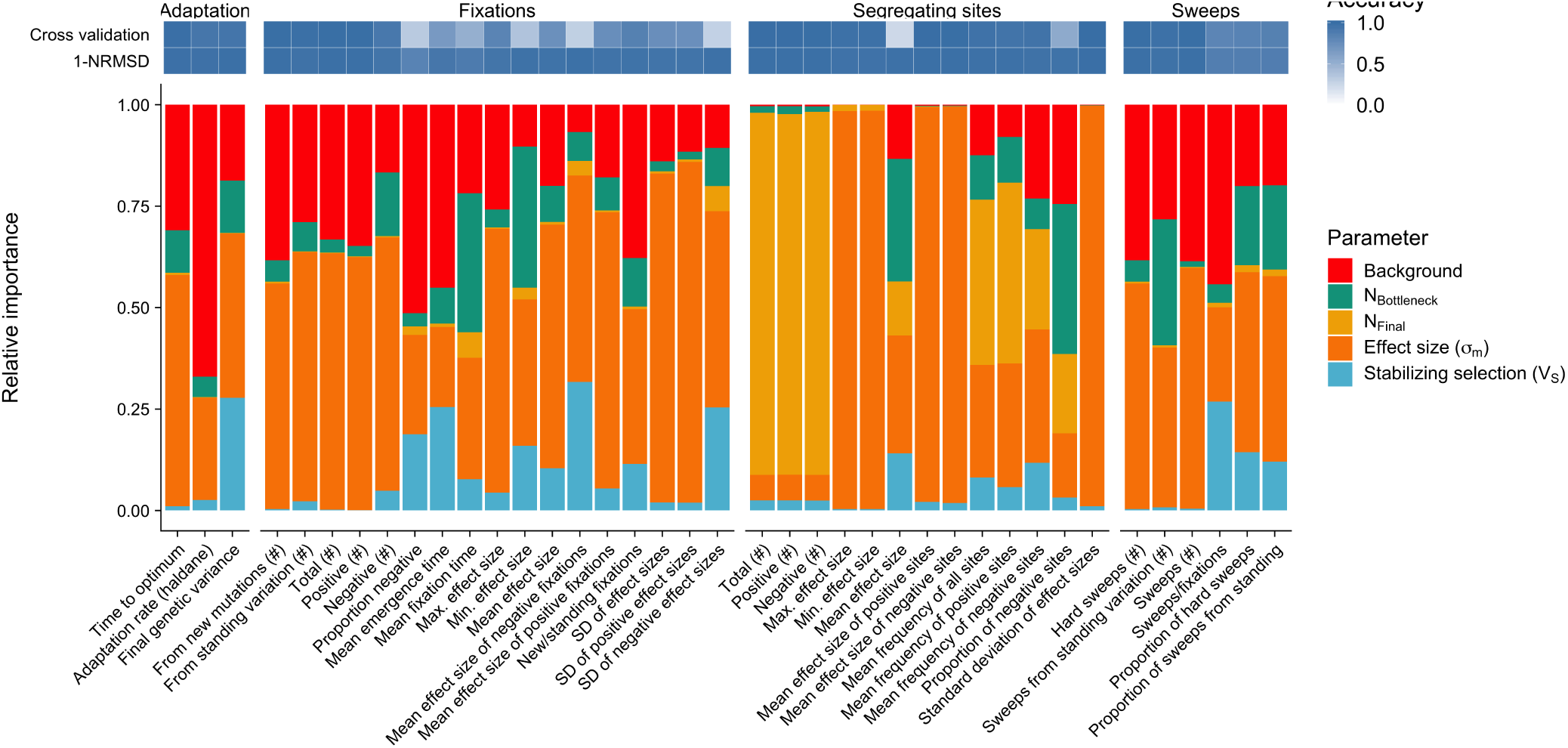
Relative parameter importance. Relative parameter importance inferred for four parameter categories. 1) Adaptation: parameters describing adaptation speed and potential for future adaptation, 2) Fixations: summary statistics for mutations that were fixed during trait adaptation, and 3) Segregating sites: descriptors of alleles polymorphic in the final generation of the simulations. Top rows indicate prediction accuracy as calculated by 10-fold cross validation and NRMSE. Each bar is the result of an independent random forest learning and each color represents the relative importance of the simulation input parameters (see Methods and Table S1 for summary statistics).

**Figure 4.**
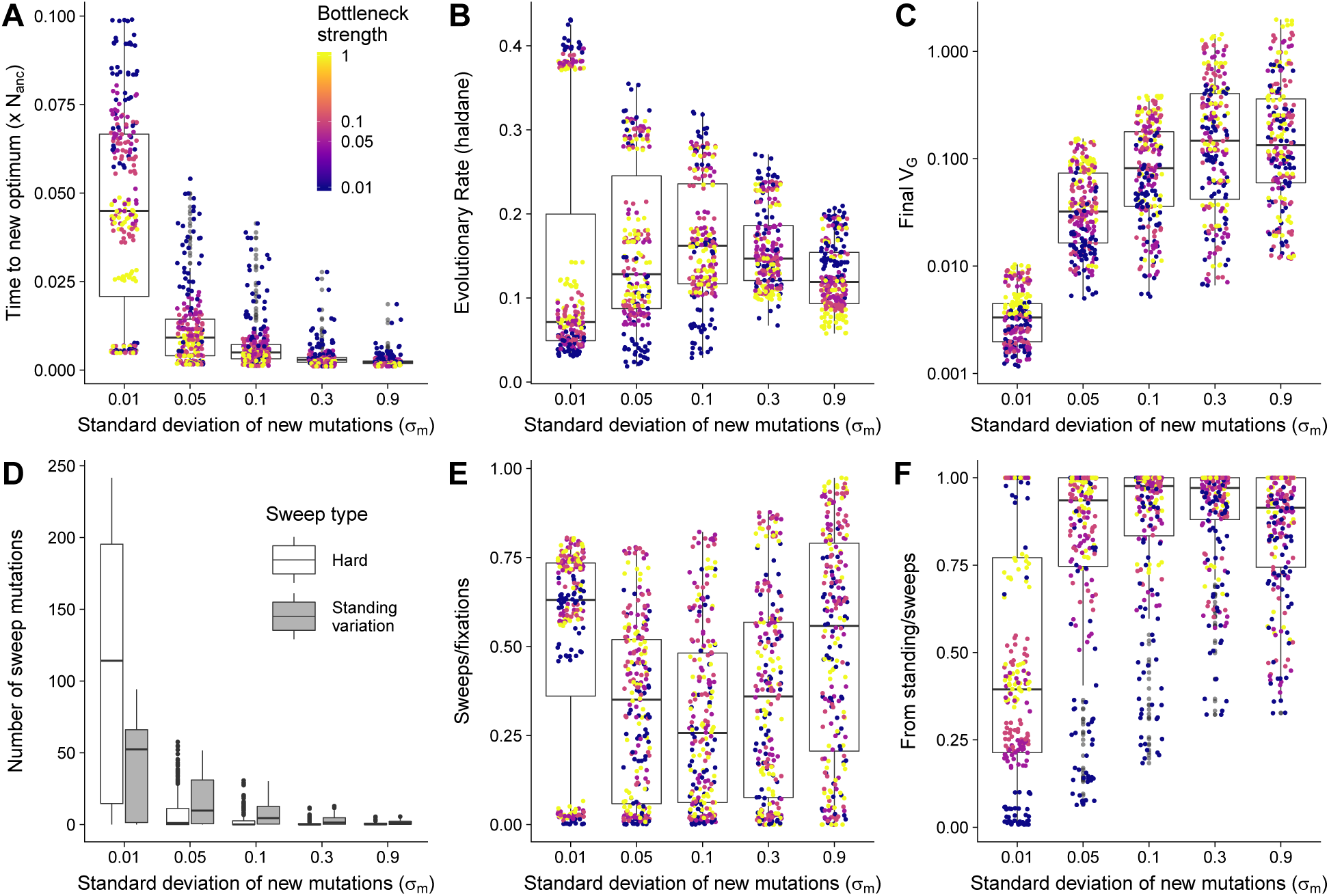
Summary of trait adaptation and selective sweeps. **A**) Time to reach new trait optimum **B**) Rate of change in phenotype **C**) Genetic variance after 0.1 × N_anc_ generations. **D**) Total number of selective sweeps, separated by type of sweeps. **E**) Proportion of sweeps compared to all fixations **F**) Proportion of sweeps from standing variation. Boxes are split by major parameter importance as identified by our random forest model. Points in A-C and E-F show the values of each of 1,200 parameter sets and are colored according to bottleneck size (Darker color indicate stronger bottleneck, see legend in A). Interactive plots are available at https://mgstetter.shinyapps.io/quantgensimAPP/

### Segregating sites after polygenic adaptation

We further investigated segregating sites in the final generation, which correspond to a modern population that has experienced an optimum shift in the past. Cross validation prediction accuracies were for most summary statistics very high (<0.9). The mean effect size of segregating sites was predicted with less accuracy, however, as all values are concentrated around zero leading to low *R*^2^values in the CV. The NRMSD, shows that the accuracy for mean effect size of segregating sites was high and that the validation data could be predicted, which allowed to infer parameter importance even with lower CV accuracy.

While absolute numbers mostly depended on the final population size, other statistics showed more distinct patterns. Allele frequencies of both negative and positive sites were strongly influenced by the demography of the population. The proportion of negative sites segregating in the population was also most strongly influenced by the strength of the bottleneck (Figure 3), but when 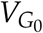 (Figure S3) was used to train the model instead, *V*_*G*_0 explained most of the variation (Figure S5). As 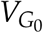 is the result of the combination of *V*_*S*_ and *σ*_*m*_ during the burn-in, this a strong interaction effect between *V*_*S*_ and *σ*_*m*_ which is partitioned when using *V*_*S*_ as feature in the random forest.

### Fixations and selective sweeps

Mutations in a population can rise in frequency and fix due to demographic events and stochastic sampling or as a result of selection. The sudden change in trait optimum in our model imposed strong selection on sites with a postive effect, while mutations with negative effect values were deleterious until the new optimum was reached. Different parameter combinations led to strongly varying numbers and patterns of fixations in our simulations. The effect size of new mutations (*σ*_*m*_) and *ψ* had the strongest influence on the absolute number of fixations and the effect size of mutations that fixed (Figure 4, 3 and S5). Variation in the mean effect size of fixations depended mostly on *σ*_*m*_, though *V*_*S*_ also contributed substantially for negative fixations. Consistent with fixations being driven primarily by selection, the effect size of positive mutations that fixed was an order of magnitude larger than that of negative fixations (Figure S6). Comparing results within each set of simulations with identical *σ*_*m*_ shows that stochastic sampling due to *N*_*bneck*_ played an important role in determining the number of fixations even if the relative importance of *N*_*bneck*_ among all parameters was only 3% (Figure S6A and 3).

Not all fixations are due to positive selection, however, and even those that are due to selection would not necessarily reduce linked diversity sufficiently to be detected as a selective sweep. To differentiate between neutral and strongly selected fixations, we compared the fixation time of sites that fixed after the shift in trait optimum to single-locus neutral simulations with identical demography (see Methods). Consistent with the higher total number of fixations exhibited, populations with smaller *σ*_*m*_ also showed a higher number of sweeps. While the maximum number of sweeps was almost 300 (for *σ*_*m*_ = 0.01, *ψ* = 0, *V*_*S*_ = 1, and a bottleneck), 13 parameter sets did not lead to any sweeps within the simulated time, all with *σ*_*m*_ *≥* 0.3, *ψ* = 0.95 and *V*_*S*_ *≥* 5. The proportion of sweeps to fixations ranged from 0 to 99% but was highly variable and revealed strong interactions between *σ*_*m*_, *ψ* and *V*_*S*_ (Figure 4). Larger *ψ* led to a low proportion of sweeps to fixations when *V*_*S*_ and *σ*_*m*_ were small, but for large values of *V*_*S*_ and *σ*_*m*_ almost all fixations were sweeps, scaling with decreasing *ψ* (https://mgstetter.shinyapps.io/quantgensimAPP/). The proportion of sweeps from standing variation was also highly variable, but differentiated more strongly by demography within each group of *σ*_*m*_ than the total proportion of sweeps (Figure 4E). Population bottlenecks were the second most important parameter for the type of selective sweep observed, while either *σ*_*m*_ or 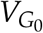 were the most important parameters 3 and S5).

### Genetic architecture after adaptation

The genetic architecture of phenotypic traits that we observe in populations today was shaped by demographic history and past selection. We evaluated the genetic architecture in the final generation of all 1,200 populations with their diverse range of histories by comparing the combined allele frequency - effect size matrices (see Methods). These frequency matrices were used as input for our random forest model to understand the contributions of input parameters to variation in genetic architecture.

The extracted parameter importance showed that the variation in the genetic architecture depended most strongly on N_final_ and *σ*_*m*_, but each of the other three parameters contributed at least 9% of the variation (Figure 5). The strong interaction between parameters becomes apparent in Figure 5, where the fine structure beyond the major 2 parameters (*σ*_*m*_ and N_final_) can be seen on all levels of combinations. Among simulations with large *σ*_*m*_ and large N_final_, however, all correlations are close to 1 and it is therefore not possible to easily distinguish parameter sets based on their genetic architecture (individual genetic architecture plots for each parameter combination are available at https://mgstetter.shinyapps.io/quantgensimAPP).

**Figure 5.**
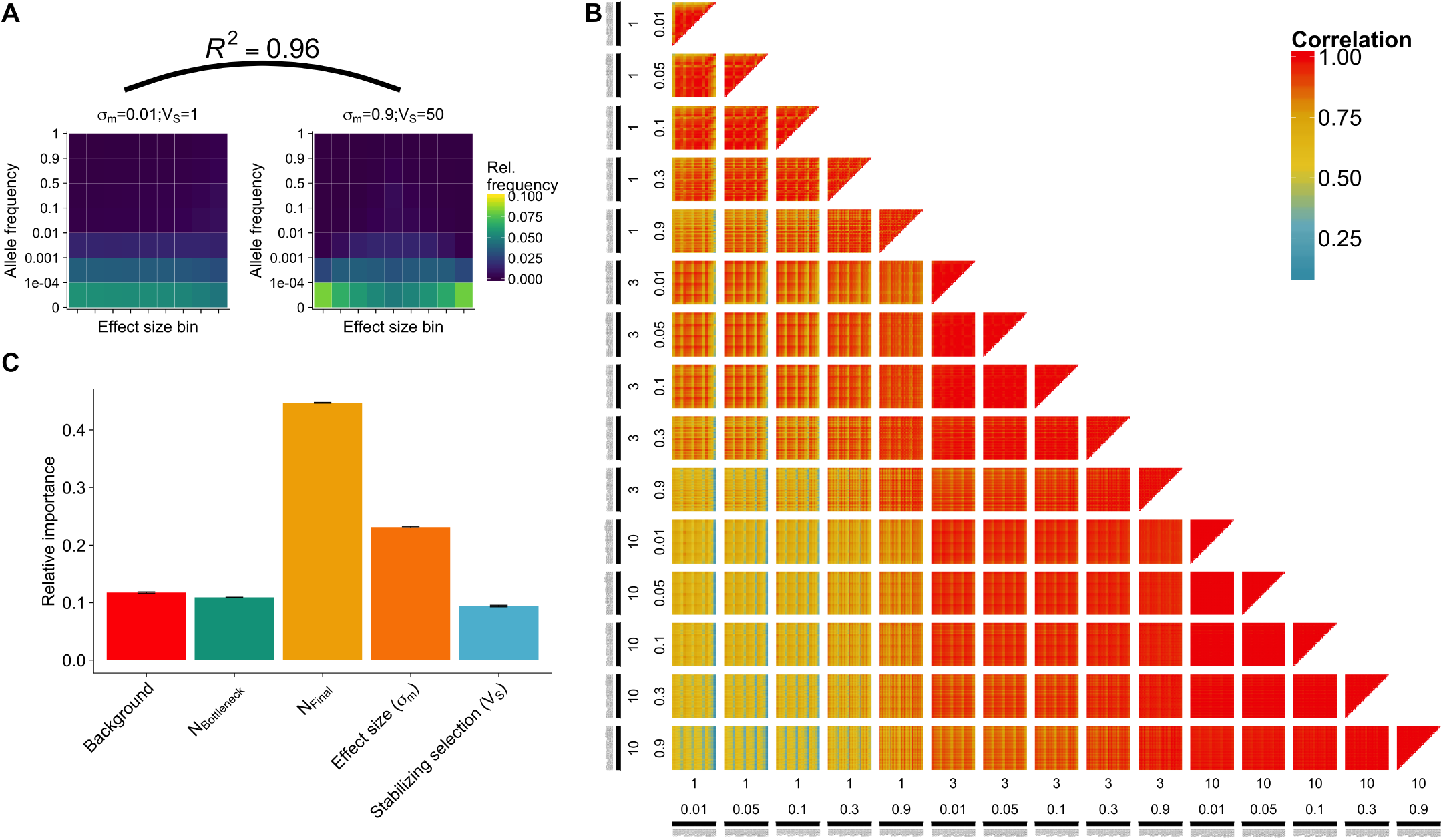
Genetic architecture in final population. **A**) Genetic architecture matrices for two parameter combinations (maize models, see Methods) differing in effect size of new mutations and strength of stabilizing selection. Effect size bins are centered around zero with negative effect size quantiles on the left and positive quantiles on the right of the central bin. Shown is the correlation coefficient between the genetic architectures. **B**) Pairwise correlation of genetic architecture of all comparisons of 1,200 parameter combinations. Subplots display the combination of final population size (log; 1, 3, 10) and effect size distribution (*σ*_*m*_, 0.01,0.05,0.1,0.3,0.9) of incoming mutations. Each pixel displays a pairwise comparison between two of the 1,200 scenarios. **C**) Relative parameter importance for genetic architecture prediction.

### Maize domestication traits

After evaluating a wide parameter space using our machine learning models, we then investigated in more detail two parameter sets that resemble diverging traits during maize domestication. Using simulations with demographic models similar to that inferred for maize [a bottleneck of 0.05 × N_anc_ followed by exponential growth to 10 × N_anc_, 22], we selected one trait with strong stabilizing selection and small effect mutations (Trait 1; *σ*_*m*_ = 0.01 and *V*_*s*_ = 1) and one trait with weak stabilizing selection and large effect mutations (Trait 2; *σ*_*m*_ = 0.9 and *V*_*s*_ = 50).

The two traits showed notably different patterns of adaptation (Figure 6, x-axis on *log*_10_ scale). Trait 1 increased almost linearly for 0.0733 × N_anc_ generations before asymptotically arriving at the new optimum. The genetic variance for this trait declined for the first 0.0169 × N_anc_ generations before it slowly increased, but did not reach the equilibrium value within the 0.1 × N_anc_ generations simulated. Trait 2, on the other hand, adapted rapidly, reaching the optimum in only 0.002 × N_anc_ generations. The genetic variance for Trait 2 increased during adaptation to a value higher than 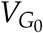, then decreased after the optimum was reached but remained higher than 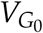 (Figure 6A and B). The number of fixations was 100 times higher for trait 1 than for trait 2; the ratio of sweeps per fixation was also higher, and most sweeps in trait 1 were hard (Figure 6C). Though on average trait 2 exhibited fewer than 2 sweeps per simulation, 94 % of these were from standing variation. Neither trait showed a strong correlation between mutation effect size and when fixation occurred, suggesting that the domestication bottleneck was not the primary driver of fixation (Figure S7). The sojourn time for sweeps from standing variation was correlated with the initial allele frequency, but also with the effect size of a mutation. Large effect positive mutations had a low initial frequency but fixed fast, while negative alleles fixed slowly, despite their high initial frequency similar to the trait we described above (Figure 2). This observation held particularly true for Trait 2, where only few small or negative effects fixed quickly (Figure 6 D and E). The overall contribution of all sweeps to phenotypic change was also different between the two traits: the summed effect size of all sweeps represents 45 % of the adaptation in trait 2, but only 18 % for trait 2.

**Figure 6.**
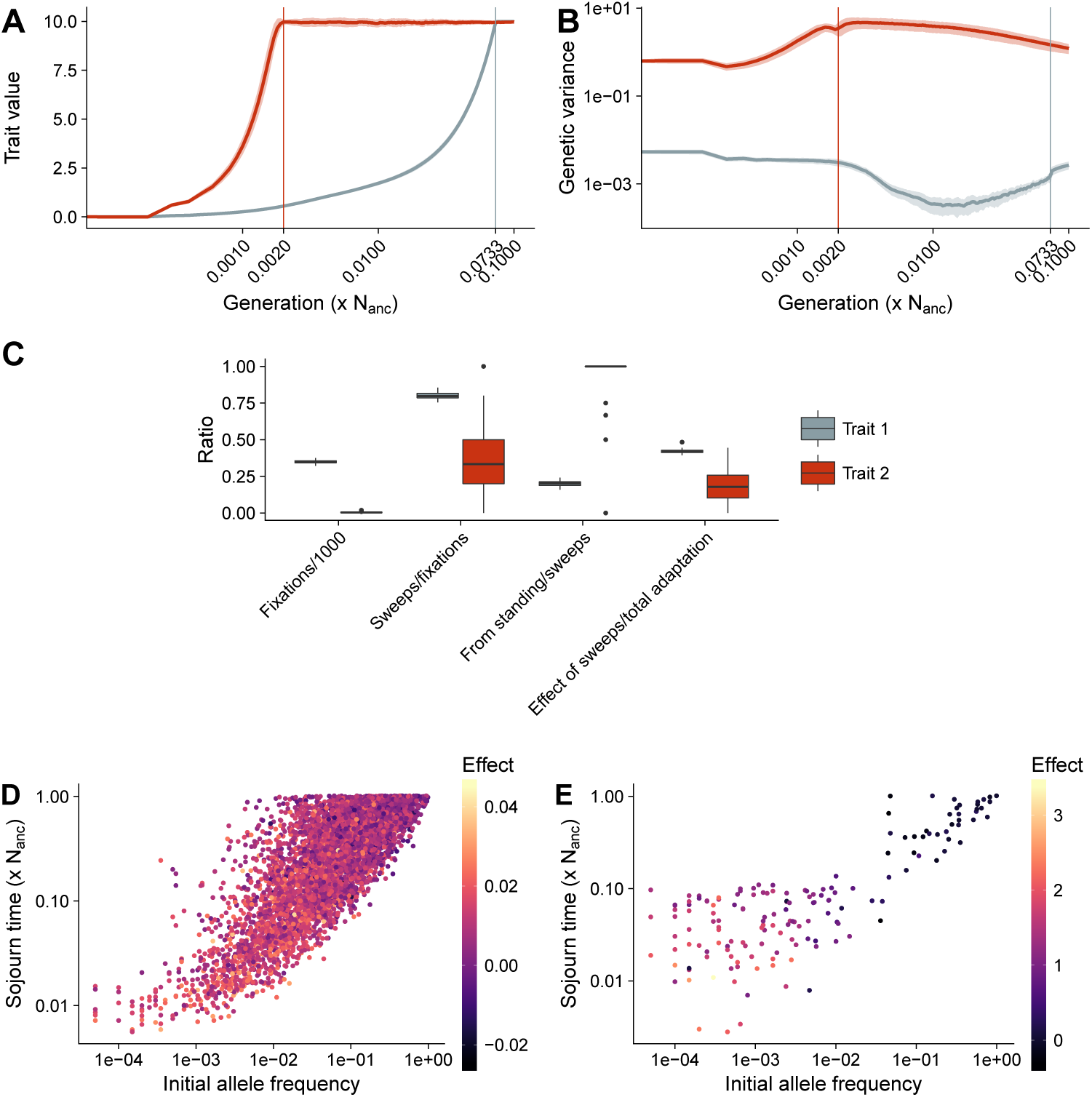
Maize specific adaptation procedure. **A**) The evolution of trait value and **B**) genetic variance during adaptation to a new trait optimum for two traits under maize demography with no genetic background. Time in both figures is shown on a log scale, light shadows show standard deviations from the mean of 100 simulation replicates. Trait 1 (blue) with small effect mutations (*σ*_*m*_ = 0.01) strong stabilizing selection (*V*_*S*_ = 1). Trait 2 (red) with large effect new mutations (*σ*_*m*_ = 0.9) and weak stabilizing selection (*V*_*S*_ = 50). Vertical lines denote the generation when 99% of the new trait optimum is reached. **C**) Proportions of selective sweeps. **D**) Sojourn time of sweeps from standing variation in Trait 1. **E**) Sojourn time of sweeps from standing variation in Trait 2. Scales in D and E are different due to strong divergence of effect size values.

Figure 5A shows the difference in genetic architecture between the two traits. While the adaptation of trait 1 led to an equal distribution of effect sizes at low frequencies, trait 2 had a larger proportion of both very low frequency mutations from the extreme tail of the distribution and small effect mutations at higher frequencies. Despite these differences the correlation between the genetic architecture matrices was very high (0.96; Figure 5).

## Discussion

### Model choice

We use a combination of two different fitness functions to study the quantitative genetics of adaptation to a sudden change to a new trait optimum far beyond observed trait values for any individual in the equilibrium population. During the stationary phase before the shift and after reaching the new optimum we followed a Gaussian fitness function appropriate for a trait under stabilizing selection [14]. During the optimum shift, however, such a model would be problematic, as only a few individuals in the upper tail of the fitness distribution would have extremely high relative fitness, inducing a strong population bottleneck. Instead, we applied a model of truncation selection, first calculating fitness under the Gaussian fitness function but then assigning a fitness of 1 to the top half of the population and 0 to the bottom half. Such a model is reasonable for sudden shifts in trait optima that do not lead to the extinction of a population, but where higher trait values are unambiguously advantageous and the maximum population size is limited. In natural populations these factors can be observed when sudden changes in the environment favor a specific phenotype for invasive species [28] or in semi-artificial populations in agroecosystems and during domestication [24]. Truncation selection is also common in evolve and re-sequence experiments [29], crop populations [30] and during strong directional selection in natural populations[31].

In our model simulations we fixed the equilibrium optimum to 0 and the new optimum to 10, but change *V*_*S*_ and *σ*_*m*_. As *V*_*S*_ and *σ*_*m*_ change, the relative distance to the new optimum changed with respect to the initial *V*_*G*_ 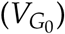. The wide range of distances simulated resembles observations in nature and experimental populations. For example, in the Illinois long term selection experiment in maize, 105 generations of selection for high oil resulted in a shift of over 40 standard deviations [30], and large trait shifts have also been identified in other experimental and natural populations [32, 33]. Our results should therefore be relevant for a variety of traits that adapt to changing environments.

While our modeling investigated a wide parameter space for a number of key variables, one key aspect we have ignored is interaction among alleles (dominance) or loci (epistasis). Both forms of interaction are widely recognized to be important at the molecular level [34, 35], but the majority of variance for a wide array of quantitative traits seems reasonably well explained under a simple additive model [36, 37], but see [38, 8, 34, 39]. And although we do not include any explicit simulation of interlocus interactions, our quantitative trait model is such that the effect of an allele in any given generation will depend on the genetic background. We predict that epistasis and dominance would absorb some of the effect of *σ*_*m*_ for most statistics and have relatively little influence on demographic parameters. Further efforts should incorporate the effects of dominance and epistasis, especially for understanding phenomena such as heterosis and inbreeding depression, where nonadditive effects are likely to play a significant role [40].

### How do organisms adapt to change

Rapidly changing environments, such as those faced by changing climate impose a threat to populations with narrow genetic variance for important traits. Quantitative traits inherently provide adaptive potential to a population, because of the genetic variance created by varying effect sizes at a number of alleles [41]. However, the speed and manner in which traits adapt depends on the initial variation and beneficial mutations entering the population once the environment changes. In rapidly changing environments or during new colonization of habitats the time it takes to reach the new optimum is critical as this might determine whether the population is first to occupy a niche. We looked at two summary statistics — time to optimum and adaptation rate — to compare the adaptive behavior of different traits. The speed to the optimum shows the absolute speed of a population to reach the new optimum, while the adaptation rate is corrected for the genetic variance present. The absolute speed depends most on *σ*_*m*_, but the adaptation rate is more uniform across *σ*_*m*_ with even higher adaptation rates for small *σ*_*m*_ (Figure 4A and B). This shows that with small effect mutations and strong stabilizing selection adaptation is mutation limited, but this is not the case when *V*_*S*_ is large. These two types of adaptation regimes have previously been described as mutation and environmentally limited adaptation regime [16]. Large adaptation rates are reached with the largest *ψ* (0.95) values, because genetic diversity is maintained during the adaptation process. Populations with small *σ*_*m*_ and small *ψ* run out of genetic variance, because most positive standing variation fixes and negative mutations get lost. The loss of genetic variance is also apparent when comparing the initial genetic variance to the final genetic variance, which is smaller after adaptation for most populations with *σ*_*m*_ = 0.01 (Figures 4 C, 6B and S3). The decrease in *V*_*G*_ for small effect mutations and the increase from large *σ*_*m*_ is consistent with previous results [15]. The genetic variance after historical adaptation is important in the face of climate change where recently adapted populations will be forced to further adapt. Populations with a large initial genetic variance and large effects also have larger genetic variance in the final population and are thus better prepared for future adaptation. The severity of population bottlenecks is an additional factor influencing *V*_*G*_ in the final generation as diversity is removed by genetic drift (Figure3 and S5).

Overall, populations with the largest 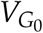 and largest *σ*_*m*_ adapt fastest to a new optimum as expected, but we also show the impact of population bottlenecks and the overlap between trait architectures (combinations of *V*_*S*_ and *σ*_*m*_). Different trait architectures can result in similar adaptation speed and genetic variance depending on the population history. This implies that for traits that are highly polygenic, it is of particular importance to prevent population declines in order to maintain the adaptability of populations.

### Selective sweeps during polygenic adaptation

Much of standard population genetic theory assumes mutations have a constant fitness effect *s*. This assumption has led to a number of findings about selective sweeps, from the probability of fixation being ≈ 2*s* [42] to the rule of thumb that mutations with fitness effects |2*Ns*| *>* 1 will be fixed or removed by natural selection, while those with smaller effects will drift stochastically as effectively neutral alleles [43]. For quantitative traits, however, the fitness effect of a mutation is conditional on the phenotypic distance of an individual to the trait optimum and the correlation between the trait and fitness [14]. At equilibrium this follows a Gaussian distribution (Equation 1), but during directional selection it will depend on the distance of the population from the trait optimum. The relationship between the phenotypic effect size of a mutation and its fitness effect is strongly positive at the onset of selection, while the slope declines as the population trait mean approaches the new optimum and is even slightly negative once the new optimum has been reached [17, Figure S8]. This shows that segregating large effect positive mutations are beneficial when the population trait mean is distant from the new optimum, but become disadvantageous once the population mean is close to the new optimum, as on average they will cause individuals to overshoot the optimum.

Most selective sweeps occur during the adaptation process before the new optimum has been reached, but the number of fixations and sweeps is strongly dependent on the demography of the population. A strong population bottleneck leads to more fixations, but most of these are fixed by drift rather then selection, and N_bottleneck_ is therefore more important for the number of fixations than the number of selective sweeps (Figure 3 and S5). Population bottlenecks also decrease the proportion of sweeps from standing variation and favor hard selective sweeps, because the bottleneck removes segregating beneficial alleles (Figure 4).

The overall importance of selective sweeps for different traits depends on the initial genetic architecture: our two example traits show that differences in the number of sweeps do not necessarily reflect their combined effect: while trait 1 exhibited 279 sweeps, these contributed to 42 % of the change in trait value, while for trait 2, only 2 sweeps contributed 22% (Figure 6C). This is consistent with previous results showing that allele frequency shifts of large effect alleles are sufficient to reach the new optimum, but selective sweeps are more important when the new optimum is distant [20, 15] Our results show even more extreme cases, for example trait 1 and simulations with *σ*_*m*_ *≤* 0.05, in which the population exhausts standing variation and relies almost entirely on new mutations. In this case hard selective sweeps are most common, as new positive mutations provide a strong relative fitness advantage (Figure 4 and 6).

Without linked neutral sites, our ability to identify likely sweep regions requires a few important caveats. First, we use a conservative definition of selective sweeps, including only those alleles fixing faster than 99% of neutral simulations. Less conservative cutoffs should not strongly influence the general result, as most mutations that sweep fix substantially faster than neutral fixations and only few more fixations would be defined as selective sweeps. Second, while we identify only sweeps from mutations that arose after the optimum shift as hard sweeps, some sweeps from standing variation would be difficult to distinguish from hard sweeps in genomic data if their frequency at the onset of directional selection was very low [44]. Likewise, not all alleles that fixed faster than 99% of neutral simulations would be detectable as selective sweeps in empirical data, as selection on standing variation has a less pronounced impact on diversity at linked sites [4].

### The effect of genetic background on focal QTL

Allele frequency shifts and selective sweeps in a focal QTL are dependent on the genetic background. [17] showed analytical results for the behavior of a single locus with a polygenic background during the adaptation to a new optimum. In our study, we simulate a more complex case: in addition to a genetic background (see equation 4), we model 20 QTL each involving numerous loci. Moreover, in our model the QTL and the genetic background are not independent, because the QTL in the parents contribute to their trait value but can themselves be inherited as well. Nonetheless, our results broadly agree with [17], showing that when the effect of the background and effects of mutations within the QTL are large, adaptation proceeds without selective sweeps (Figure 4). We additionally show that the background explains considerable variation in many summary statistics, in particular those related to fixations and selective sweeps (Figure 3). Together with empirical observations of varying fitness effects for QTL in different backgrounds [45, 46, 47], our results highlight that evolutionary models of QTL cannot ignore the effects of genetic background.

### Genetic architecture of quantitative traits after adaptation

The genetic architecture of a trait is an important feature in the study of adaptation, influencing both the response to selection as well as the power to detect causal loci for a trait. Our two example traits show that different adaptation processes lead to different patterns of the genetic architecture matrix. Because trait 1 only reached the new optimum shortly before we assess the the genetic architecture, the values are distributed asymmetrically along the zero effect size bin. Trait 2 reached the new optimum very early and therefore is more similar to an equilibrium genetic architecture with effects sizes close to zero at higher frequency and larger effect sizes at very low frequencies. These differences even between two highly correlated genetic architectures show that in addition to the input parameters, the time that passed since the new optimum was reached has an influence on the genetic architecture we observe in a population.

Using a machine-learning approach that trained on a subset of our simulations, we were able to identify the parameters that explained the largest proportion of variation among the genetic architectures studied (Figure 5). We found that demographic change plays a key role in determining the present genetic architecture, explaining as much as 55% (growth and bottleneck combined) of the variation we observed. For example, recent population growth leads to an increased number of low frequency mutations; this effect drives many of the observed differences between genetic architecture matrices of different demographies. We observed a high correlation (0.83 – 0.99) between genetic architectures with similar population demographies, suggesting that making inference about the process of adaptation from present day genetic architecture will have greater power in situations where the demography can be independently inferred. The result confirms the theoretical prediction that the combination of different allele frequency shifts at a large number of loci lead to similar trait architecture [48]. However when other statistics, such as information about fixations, effect size distributions observed in present populations, number and type of selective sweeps and the demography are added as parameters to the modern genetic architecture, we can predict the evolutionary rate, *σ*_*m*_, and *V*_*S*_ with 70% accuracy.

In addition to the effect of population growth, other input parameters do contribute substantially to variation in the genetic architecture, including the strength of stabilizing selection. [49] and [50] suggest that rare alleles are unlikely to contribute substantially to trait variance, but our models show that rare alleles can explain a large proportion of the variation when effect sizes are large. This is more consistent with the findings of [21], who showed that population growth leads to an increase proportion of genetic variance explained by rare alleles. The lack of consensus might result from differences in the models: while [49] models selection on fitness directly and [50] a quantitative trait under stabilizing selection with pleiotropy, our models and that of [21] consider selection on traits that are directly correlated to fitness.

### Maize domestication

Quantitative traits have been extensively studied in maize and breeders have made steady progress selecting traits for ever increasing trait values. But despite decades of observation that many important traits in maize are polygenic and work identifying QTL underlying domestication-related phenotypes [51], there has been little attention to the process of quantitative trait adaptation during maize domestication [but see 52, 23]. Many domestication traits, alike maize traits, are polygenic and controlled by a number of loci with varying effect sizes [23]. Archaeological records of maize domestication traits show that adaptation took several thousand years [24]. Our example trait 1 matches this pattern, representing an adaptation time of almost 0.1N generations, equivalent to 10,000 years for a population similar to that of the wild ancestor of maize [22, Figure 6]. Trait 1 also leads to a reduction in genetic variance compared to the equilibrium population (wild ancestor), again matching observed data [23].

Trait 2, on the other hand, differs dramatically in a number of ways. It reached the new optimum extremely quickly, and diversity in the present is actually slightly higher than at the time of the optimum shift (Figure 6). The behavior of trait 2 most closely resembles that of resistance traits with few large effect QTL potentially [53]. We only look at the genetic variance of mutations that affect a single trait, the overall diversity of a population is based on a combination of traits with different trait architectures and neutral parts of the genome. The reduction in diversity could partly be due to the distant optimum shift and partly because of the population bottleneck experienced during domestication.

The difference in trait adaptation and genetic variance trajectory can be partially explained by the fixations and selective sweeps of beneficial alleles. The number of fixations revealed that as expected far more mutations fixed for trait 1 than for trait 2, as in trait 1 much more sites are segregating in the equilibrium population, but the number of sweeps was also much higher. This is corrected for sites that fix due to genetic drift and shows that the larger relative distance to the new optimum changes the pattern. In maize it has been shown that the domestication led to an accumulation of deleterious alleles, which so far was mainly attributed to the domestication bottleneck because no increase in deleterious alleles near major domestication genes was found [54]. For quantitative traits the small deleterious fixations could be distributed more uniformly across the genome and fix even without population bottlenecks. In general there are only few hard selective sweeps observed in maize and 84% of fitness related SNPs were already segregating in teosinte [55]. Our traits show that depending on the relative distance to the new optimum the type of selective sweeps changes. While sweeps come primarily from standing variation for traits that are close to the new optimum, for distant optima hard sweeps are observed more frequently because standing variation is exhausted. The overall pattern of selective sweeps in the maize genome is a result of selection on a combination of traits and probably involves pleiotropic effects that can prevent fixation of new mutations even if they have large effects on a single trait. [47].

### Signature of polygenic adaptation in genomic data

The recently suggested omnigenic model predicts that regulatory networks are sufficiently interconnected that many loci even outside the most biologically relevant genetic pathways can nonetheless affect a trait [56]. If indeed many traits are omnigentic, a quantitative evolutionary model as employed in our simulations is well suited for making inferences about observations in genomic data. Large sets of genomic and phenotypic data are becoming increasingly available, facilitating the study of the role of polygenic adaptation. Our results help to understand the implications of different theoretical parameters for the interpretation of such studies and provide targets for new selection tests that explicitly test for polygenic adaptation and the underlying genetic architecture. We show, for example, that selective sweeps can have a crucial role during polygenic adaptation and should be integrated into detection methods, as some approaches to investigate polygenic adaptation from shifts in allele frequencies may lose power if large effect alleles are fixed in the population in which effects are estimated [6, 39, 57].

Inferring polygenic adaptation and the underlying parameters in empirical data can provide important insight into the evolution of complex phenotypes. For experimental evolution scenarios in which the ancestral populations are known, the distance between the initial and the final optimum can be inferred from phenotype data, but for natural populations this may be more challenging. Our results indicate that the relative distance could be inferred from genomic data via estimates of the genetic architecture if the demographic history is known. One current challenge of transferring simulation results to empirical data is the computational limitation of simulating whole genome sequences in large populations. Faster implementations will allow simulation of larger regions and include neutral sites [58], and could be used to train machine learning models in order to predict the evolutionary history of a population from existing data coming from association studies. The implementation of machine learning trained on simulated data has been successfully applied to identify a number of population genetic patterns [59], and is a promising avenue for future work.

## Materials and Methods

### Model

We simulated a quantitative trait under stabilizing selection with an optimum of 0 that adapted to a discrete optimum change to a value of 10. The population was diploid and mated randomly. Phenotypes followed a purely additive model in which the genotypic values at a given locus with an allele of effect size *a* were 0, 0.5*a* and *a* for homozygous ancestral, heterozygous and homozygous derived genotypes. We modeled 20 QTL resembling 50kb regions, each with a 4 kb “genic” region centered in a 46 kb “intergenic” region. In the intergenic region mutations that affect the phenotype appeared with 1% probability of the genic region, leading to approximately 10% of mutations in intergenic regions and 90% in the 4kb genic regions. Starting with a neutral substitution rate of 3 × 10^-8^per site per generation [60], we then assumed that only 10% of all mutations affect the trait of interest, resulting in a mutation rate of 3 × 10^-9^per site per generation and a total per gamete mutation rate of 3 × 10^-3^per generation. Regions were unlinked (50 cM distance), and within regions the recombination rate was 5 × 10^-8^per site per generation (0.5 per gamete).

***Fitness*** We used a Gaussian fitness function in which an individual’s fitness *w* was modeled as:

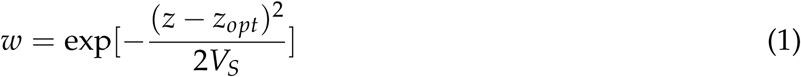

where *z* is the trait value of an individual, *z*_*opt*_ is the population optimum trait value and *V*_*s*_ modulates the possible deviation from the optimum. This standard model for traits under stabilizing selection is well suited for populations at equilibrium [26, chapter 7]. Under strong directional selection, however, this model greatly amplifies fitness differences among individuals in the tails of the phenotypic distribution. During the adaptive phase of the simulation, we calculated individual fitness following equation 1, but then apply truncation selection by assigning a fitness of 1 to the top 50% of the distribution of *w* and 0 for the remaining 50%. This model allowed for truncation selection on *z*, while the population was distant from the new optimum, but allows for selection against phenotypes that surpass the new optimum during the final stages of adaptation. We stopped truncation selection once the population mean reached the new optimum, returning the population to stabilizing selection using fitness values calculated in equation 1.

***Initial genetic Variance*** The genetic variance at equilibrium can be approximated by the house of cards (HoC) approximation [12, 26]:

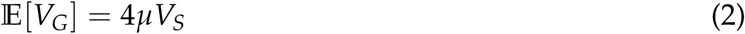

and the stochastic HoC approximation [chapter 7 26, 27]:

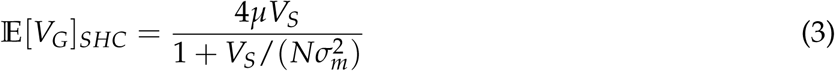

We simulated five different values of *V*_*S*_ (1, 5, 10, 20, 50) to modulate the genetic variance of the equilibrium population.

***Effect size of new mutations*** We used a Gaussian distribution around zero for the effect size of new mutations and five different standard deviations (*σ*_*m*_ = 0.01, 0.05, 0.1, 0.3, 0.9) to create traits with different effect sizes. Given a fixed optimum of 10, this distribution of effect sizes in combination with *V*_*S*_ effectively parameterize the distance to the new optimum, from a minimum distance of 11.5 z-scores (phenotypic standard deviations) to a maximum of 158.2 z-scores.

***Background*** Computational limitations do not allow simulation of an entire eukaryotic genome, so we added a heritable background (*G*_*B*_) to our simulations to account for the adaptive potential of the rest of the genome.

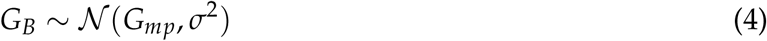

where *G*_*B*_ is the value of the genomic background of an individual, *G*_*mp*_ is the mid-parent genotypic value and *σ*^2^ is the variance of the parental trait values [61, chapter 9]. Hence, *G*_*B*_ is drawn from a normal distribution around the mid-parent value.

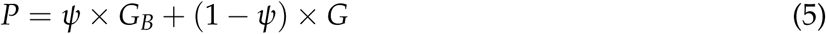

The trait value of an individual *P* is then given by the sum of its genetic value *G* and the genomic background *G*_*B*_, weighted by *ψ*, the proportion of trait variation represented by background. We modeled four different background levels (*ψ* = 0, 0.1, 0.5, 0.95).

***Demography*** To study the effect of population bottlenecks and expansion, we simulated a total of 12 different demographic scenarios with varying strength of a single bottleneck and subsequent growth (Figure S1). In scenarios with a bottleneck, an instantaneous reduction in population size occurs immediately after the burn-in and is followed by exponential growth over the length of the simulation (0.1 ×*N*_*anc*_ generations).

### Simulations

Using the above described parameters we simulated 100 replicates each of 25 different equilibrium traits using fwdpy11 v1.2a (https://github.com/molpopgen/fwdpy11), a Python package using the fwdpp library [62]. These 25 traits differed in their combination of *V*_*S*_ and *σ*_*m*_ and were run for a burn-in of 10 N_anc_ generations (Figure S3). Subsequently, each of the 1,200 parameter combinations was run for 0.1 N_anc_ starting from these equilibrium traits.

We simulate a population of 10,000 individuals for 1,000 (0.1N_anc_) generations after a burn-in of 100,000 generations to reach equilibrium.

The population mean trait values and variances were recorded every generation and entire populations, including individual trait values, mutations and effect sizes, were recorded every 10 generations for the first 100 generations after burn-in and then every 100 generations thereafter.

### Analysis

***Sweeps*** To identify selective sweeps, we used binomial sampling to simulate the sojourn time of neutral alleles arising in populations undergoing each of the demographic models described above. Mutations that were lost or that fixed before the end of the burn-in were ignored. We ran 10,000 replicates for each of the 12 demographic models and recorded the time it took a mutation that fixed within the last 0.1*N* generations (similar to our selection model) to fix in this random model. These simulations provided a null distribution to which we compared selected mutations in our quantitative trait simulation (Figure S2). We defined as a sweep any mutation that fixed faster than 99% of neutral alleles and categorized them as hard or from standing variation depending on whether the mutation arose before or after the optimum shift.

***Machine learning*** For each of the 120,000 simulations we calculated various summary statistics using the pandas version 0.21.0 and numpy version 1.12.1 Python libraries [63, 64]. These include statistics related to adaptation, selective sweeps, segregating sites, and fixed mutations; Table S1 contains a full list of parameters used for prediction and importance estimation.

To identify the importance of input variables we trained a random forest and extracted the relative importance of the input parameters. We employed the RandomForestRegressior of sklearn 0.19.0 [65] with 100 trees to extract parameter importance by training the model using input parameters as features of the whole dataset and predicting a summary statistic. The prediction accuracy for all parameters was then estimated by 10-fold nested cross validation (training using 80% of the data) as well as root-mean-square deviation normalized by the range of values observed (NRMSD) implemented in sklearn [65], and the process repeated for each summary statistic of interest (Table S1).

To compare the genetic basis of traits between scenarios we define the genetic architecture as the matrix of allele frequencies and effect sizes for each simulation. Allele frequencies were split into 7 discrete bins (0 - 10^-4^, 10^-4^- 10^-3^, 10^-3^ - 10^-2^,10^-2^ - 0.1, 0.1 - 0.5, 0.5 - 0.9, 0.9 - 1) and effect sizes were split into 9 quantiles, as absolute effect sizes were strongly dependent on the input effect size. Relative occurrence frequencies (summing to 1 over the whole matrix) of segregating sites in each frequency-effect size combination were calculated for each simulation. These values were used to train a random forest model and extract parameter importance. Parameter importance was estimated by predicting frequencies of each effect size bin from the input parameters. Prediction accuracy was again assessed by 10-fold cross validation implemented in the sklearn [65]. Additionally, we calculated pairwise correlations of genetic architecture matrices in the final generation between all possible pairs of scenarios using the mean of all simulation replicates.

***Maize domestication***

We took a closer look at two sets of simulations that represent diverging traits under a demographic model similar to that of maize domestication (N_bottleneck_ = 0.05 × N_anc_; N_final_ = 10 × N_anc_). For these simulations we assumed no genetic background (*ψ* = 0). Trait 1 represents a trait with new mutations of small effect (*σ*_*m*_ = 0.01) and strong stabilizing selection (*V*_*S*_ = 1), while Trait 2 has new mutations of large effect (*σ*_*m*_ = 0.9) and weaker stabilizing selection (*V*_*S*_ = 50).

### Data availability

All scripts and code to reproduce the simulations and figures is available at https://dx.doi.org/10.6084/m9.figshare.6179219. A detailed interactive graphical analysis of summary statistics is available at https://mgstetter.shinyapps.io/quantgensimAPP/

## Acknowledgments

We acknowledge the financial support of NSF Plant Genome award IOS-1238014, USDA Hatch project CA-D-PLS-2066-H to JRI and the Deutsche Forschungsgemeinschaft (DFG) grant STE 2654/1-1 to MGS. We would like to thank Patrick Thorwarth and Bernd Stetter for recommendations on machine learning, and CSI Davis, Graham Coop and Michael Turelli for helpful discussion. We are also grateful for the “wes anderson” color palettes that we employed for plots in this manuscript (https://github.com/karthik/wesanderson).

## Supplement

**Figure S1.**
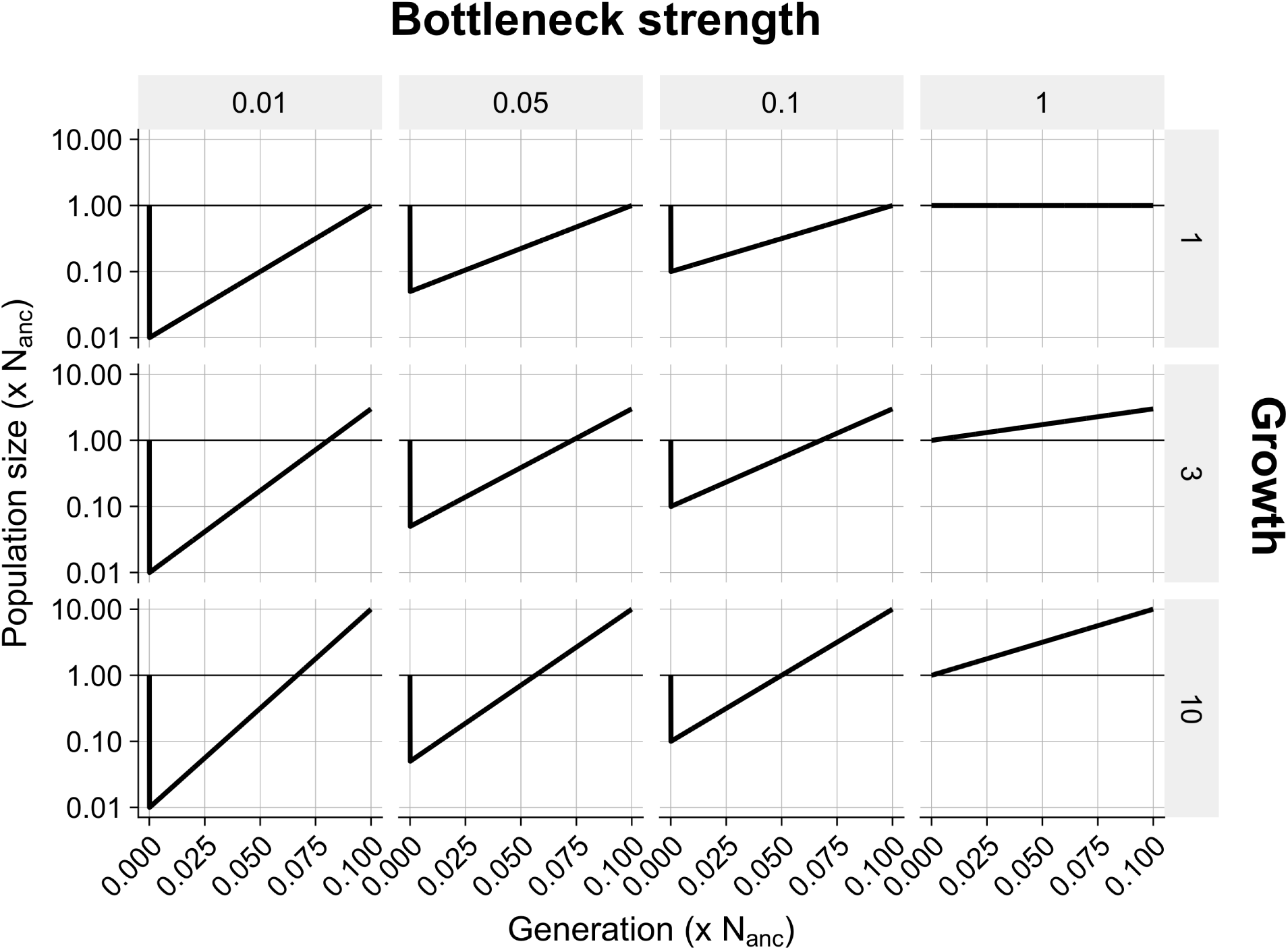
Demographies. Bottlenecks and growth models

**Figure S2.**
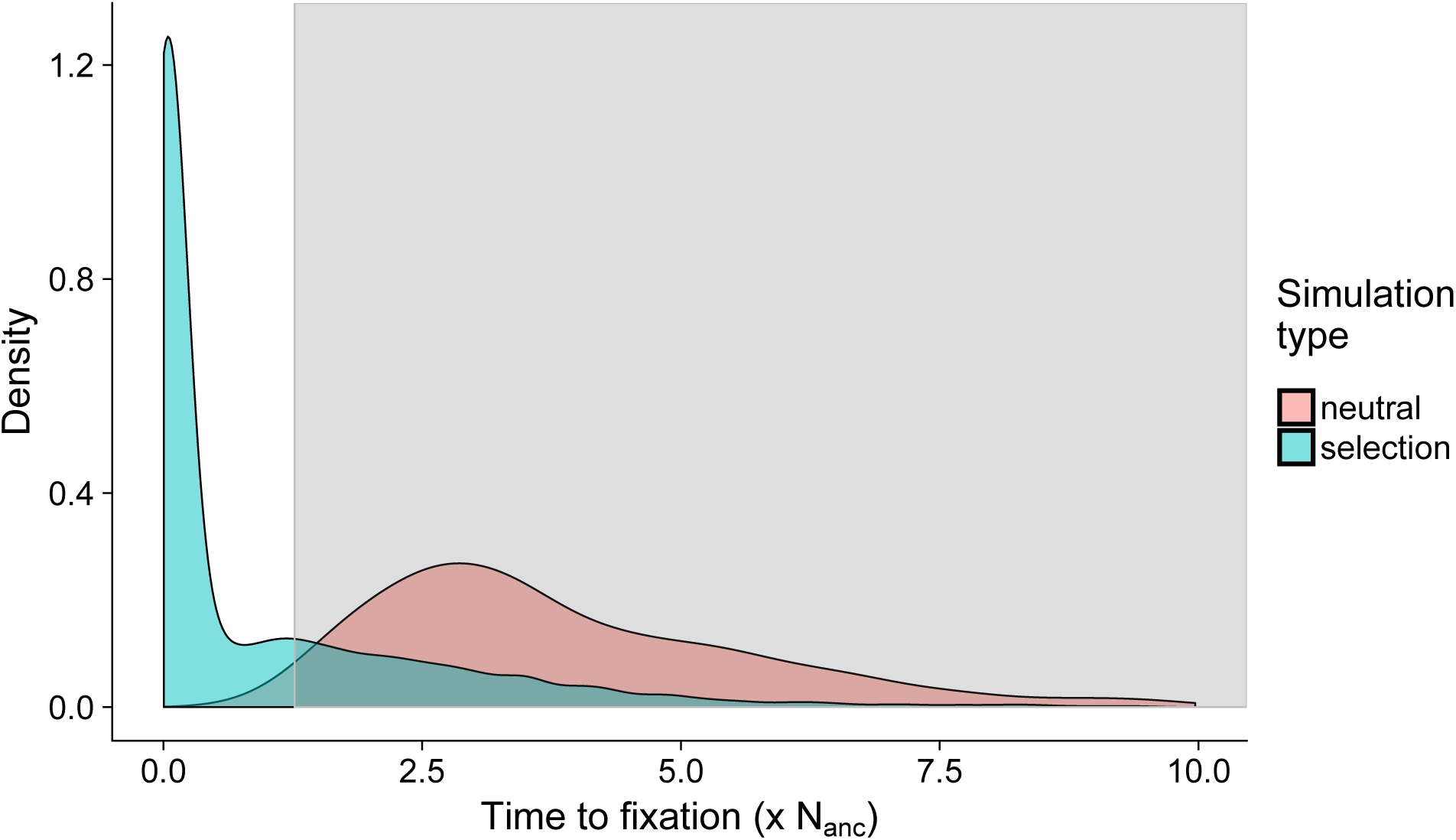
Detection of selective sweeps. Distribution of fixation times from neutral single locus simulations (red) and forward simulations with selection (green). The grey area denotes the 99% confidence interval of neutral fixation time. Fixations outside the confidence interval are considered selective sweeps.

**Figure S3.**
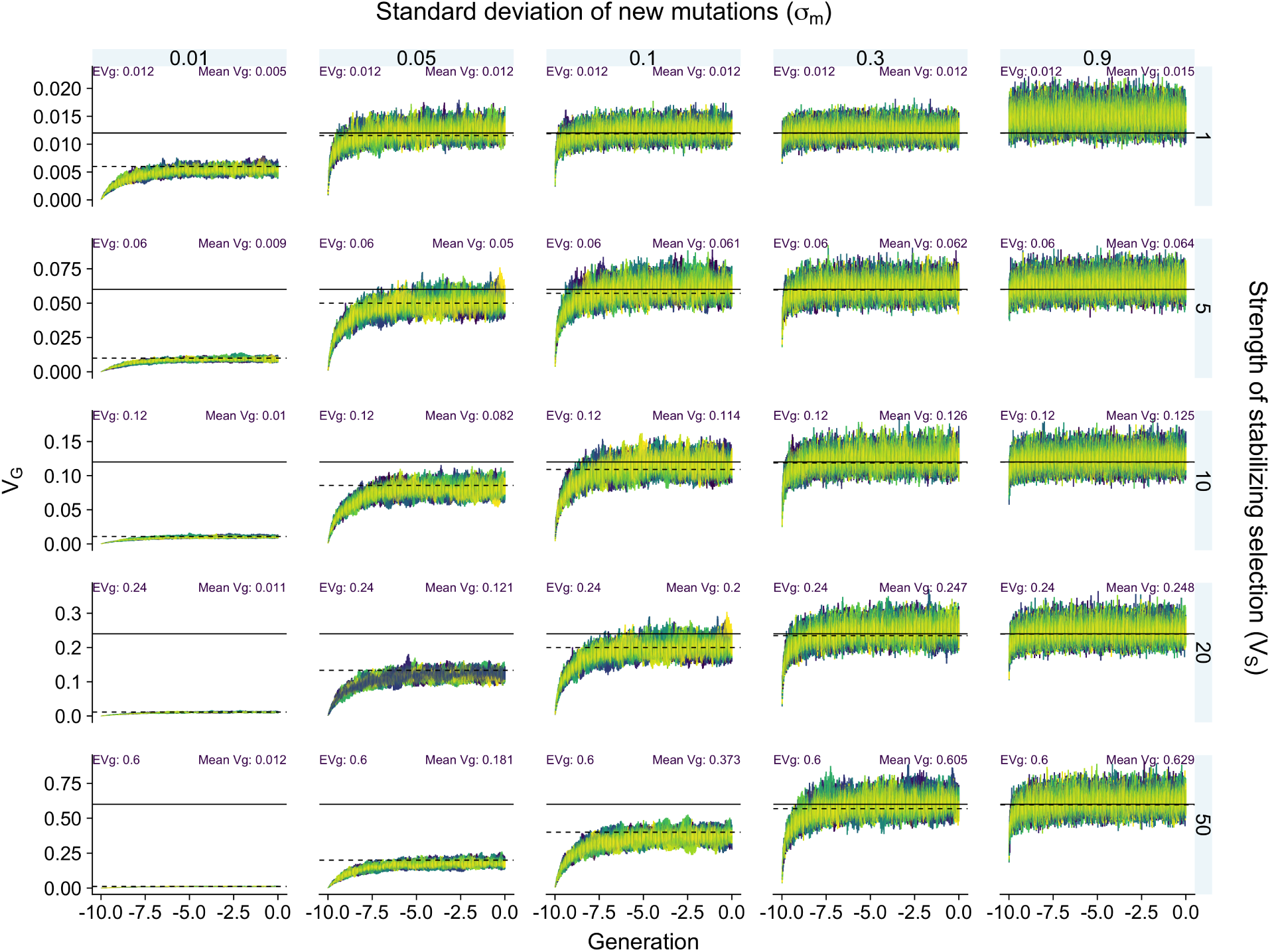
Genetic variance during burn-in. The genetic variance in each generation over 10 N_anc_ generations for each parameter set. Solid horizontal lines denote the House of Cards approximation of *V*_*G*_ [12]. Scenarios with small *σ*_*m*_ and large *V*_*S*_ do not reach the expected *V*_*G*_ because mutations are too small to “fill up” the variance volume. However, their equilibrium variance is well approximated by the stochastic House of Cards approximation [27, dashed line]

**Figure S4.**
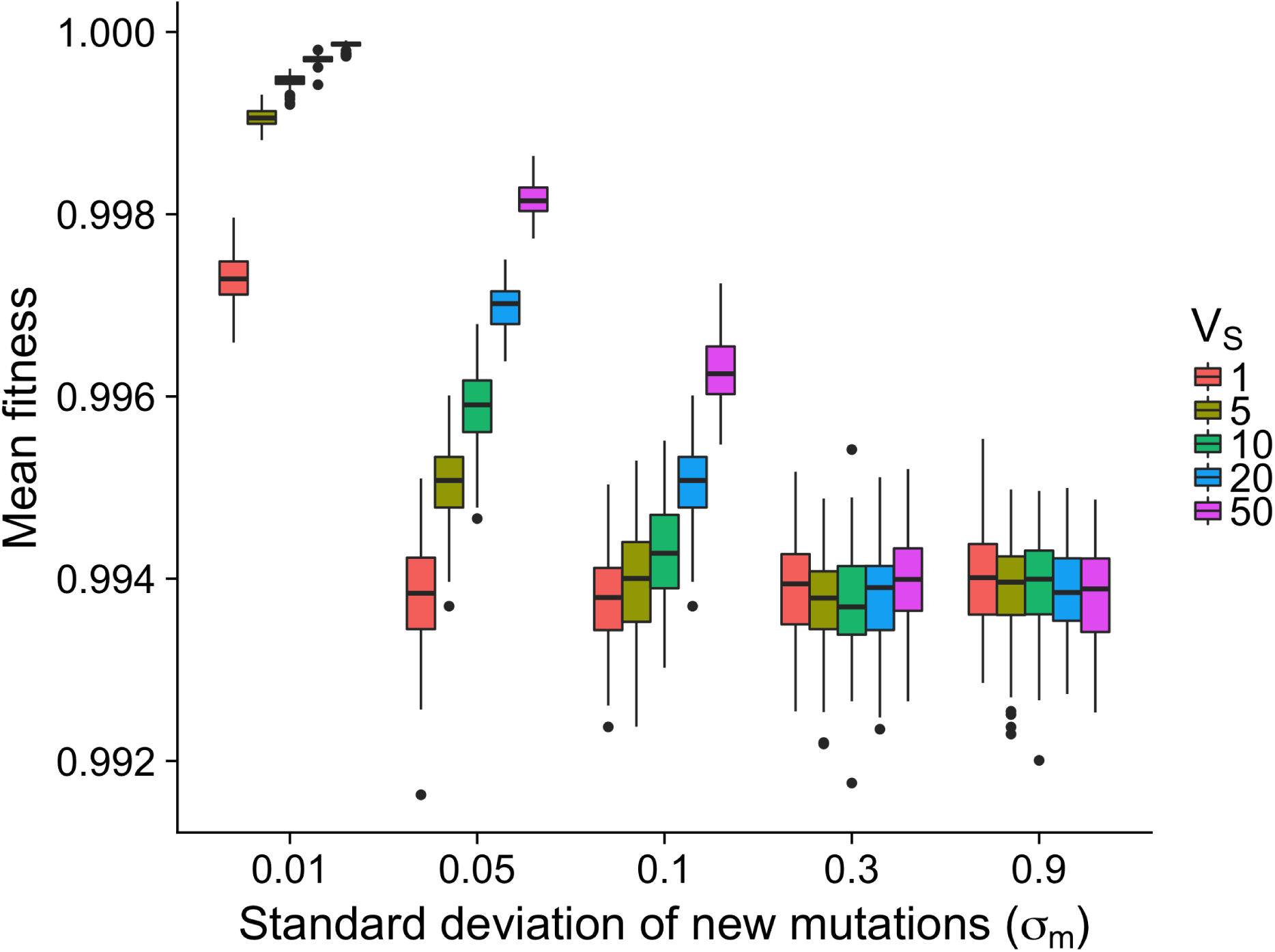
Equilibrium Fitness. Fitness for each burn-in parameter combination after 10N generations.

**Figure S5.**
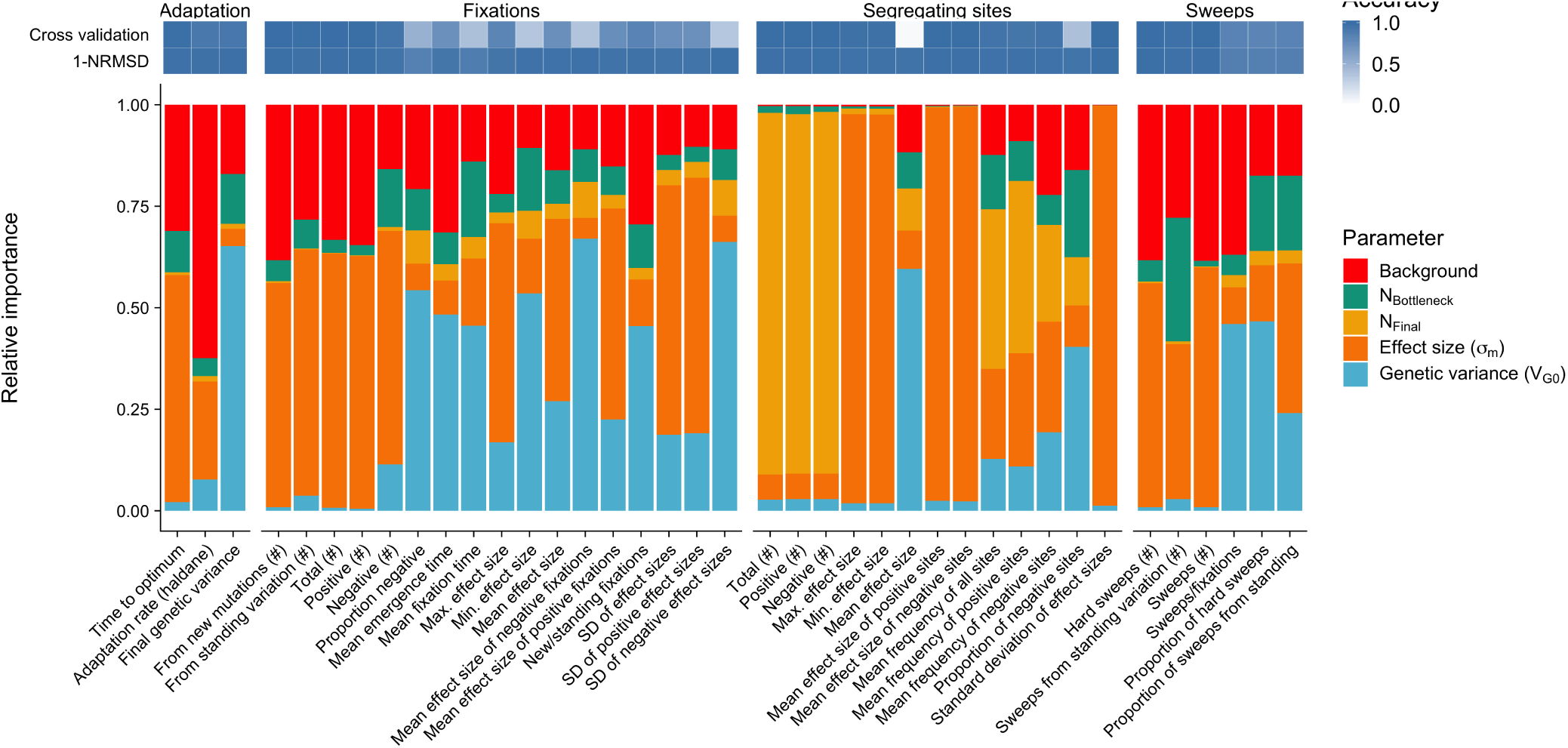
Relative parameter importance. Relative parameter importance inferred by Random Forrest machine learning for three parameter categories. 1) Adaptation, trait related parameters describing adaptation speed and potential for future adaptation. 2) Fixations, summary statistics for mutations that were fixed during trait adaptation and 3) segregating sites in the final generation of the simulations. Top panel indicating prediction accuracy as calculated by 10-fold nested cross validation and normalized relative mean squared error

**Figure S6.**
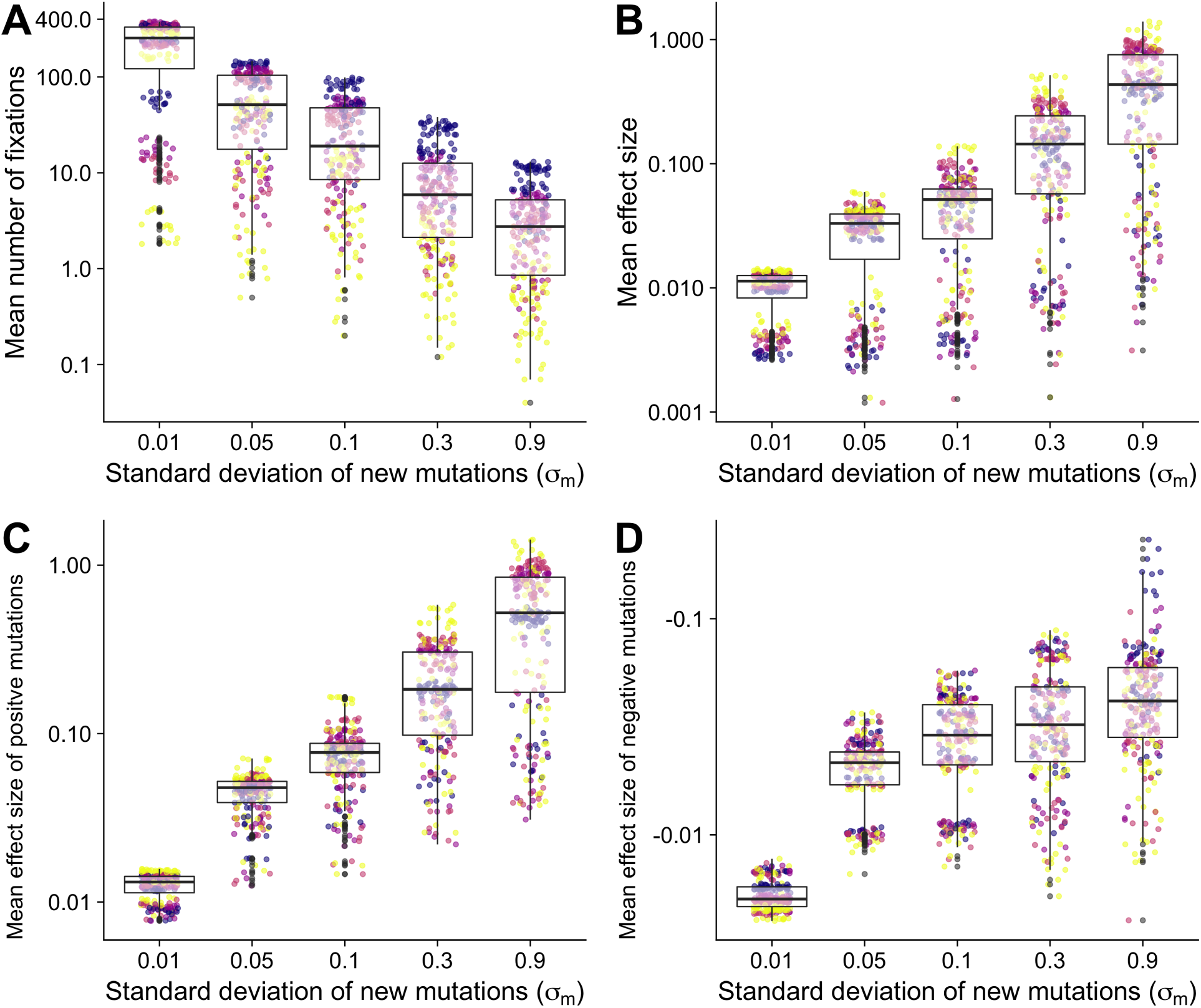
Fixations. **A**) Total number of fixations **B**) Mean effect size of fixations **C**) Mean effect size of positive fixations **D**) Mean effect size of negative fixations

**Figure S7.**
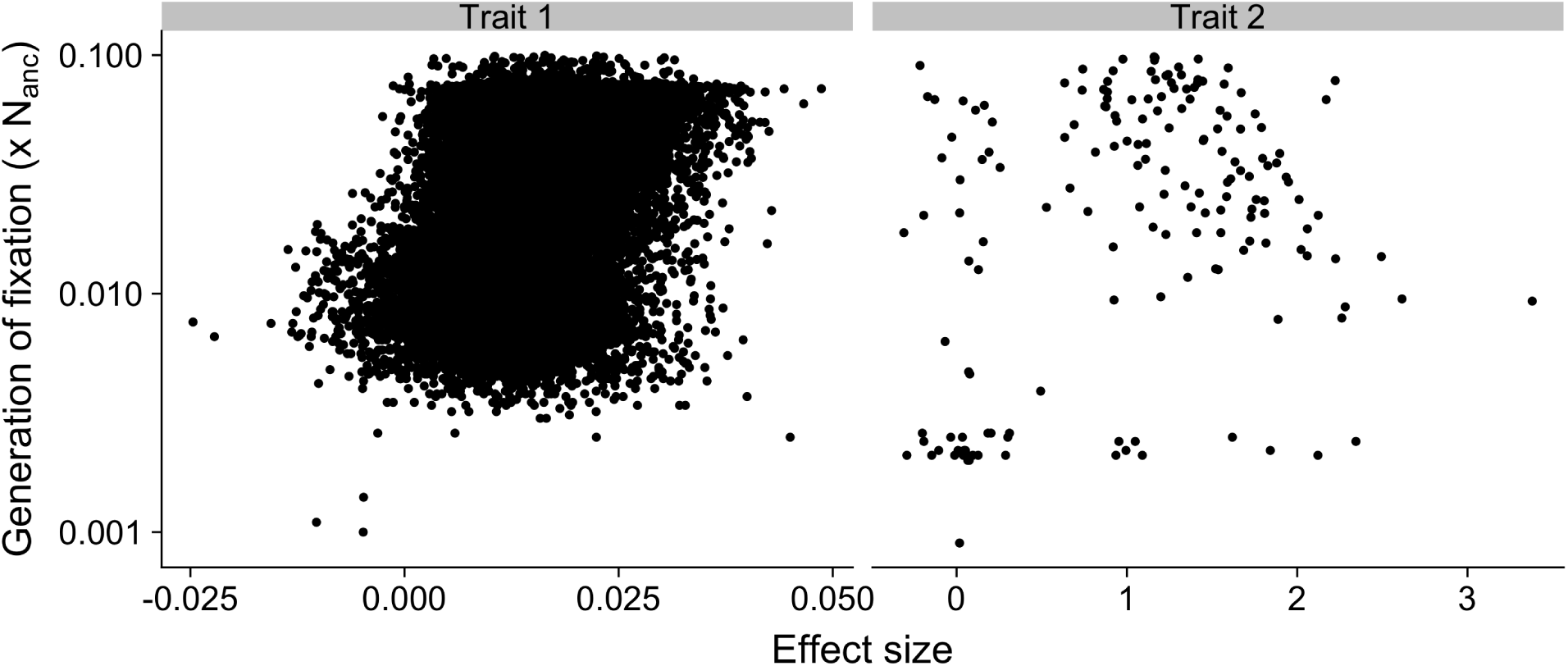
Timing of selective sweeps for maize domestication simulations. Shown is the relationship between effect size and the generation of fixation for mutations for Trait 1 (left, *σ*_*m*_ = 0.01 and *V*_*S*_ = 1) and Trait 2 (right, *σ*_*m*_ = 0.9 and *V*_*S*_ = 50).

**Figure S8.**
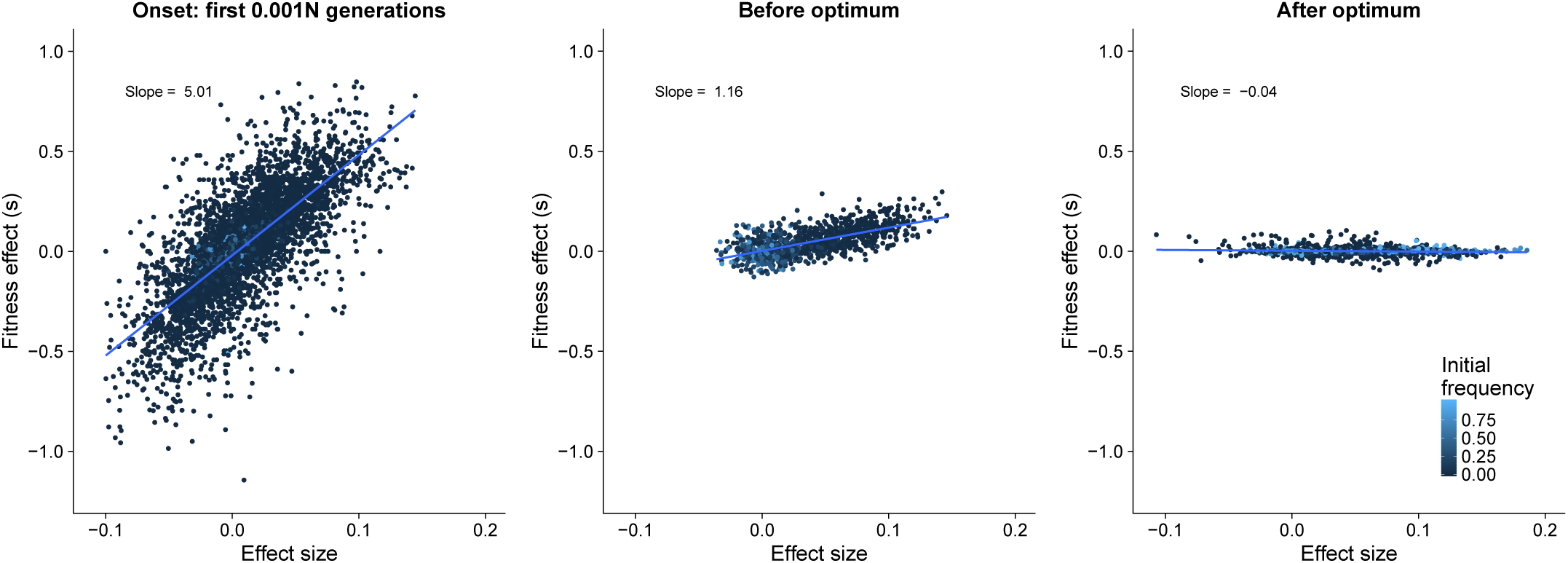
Fitness effect of mutations. Fitness effect of mutations at the onset of directional selection (0.001-0.012N), Before the new optimum is reached (0.001 - 0.012N) and after the new optimum has been reached (0.012 - 0.022N)

**Table S1.** Predicted summary statistics for feature importance estimation.

| Parameter | Description |
| --- | --- |
| **Adaptation** | **Trait related parameters** |
| Time to optimum | Generations until new optimum is reached |
| Adaptation rate (haldane) | Adaptation rate until new optimum is reached. Calculated as $rate(h) = \frac{\frac{\ln(x_2)}{sd_{x12}} - \frac{\ln(x_1)}{sd_{x12}}}{t_2 - t_1}$ |
| Final genetic variance | Genetic variance in the final generation |
| **Fixations** | **Mutations that fix after the optimum shift** |
| From new mutations (#) | Sum of fixed mutations in the final population that were already segregating before the optimum shift |
| From standing variation (#) | Sum of fixed mutations in the final population that arose after the optimum shift |
| Max. effect size | Maximal effect size of all fixations |
| Mean effect size | Mean effect size of all fixations |
| Mean effect size of negative fixations | Mean effect size of negative mutations |
| Mean effect size of positive fixations | Mean effect size of positive mutations |
| Mean emergence time | Mean generation when a mutation arose that fixed in the last 0.1 N generations |
| Mean fixation time | Mean generation in which a mutation fixed |
| Min. effect size | Minimal effect size of all fixations |
| Negative (#) | Sum of fixed mutations with negative effects in the final population |
| New/standing fixations | Ratio of mutations from new mutations vs. standing mutations |
| Proportion negative | Proportion of negative fixations from all mutations |
| Positive (#) | Sum of fixed mutations with positive effects in the final population |
| SD of effect sizes | Standard deviation of effect sizes of all fixations |
| SD of negative effect sizes | Standard deviation of effect sizes of negative fixations |
| SD of positive effect sizes | Standard deviation of effect sizes of positive fixations |
| Total (#) | Sum of fixed mutations in the final population |
| **Sweeps** | **Mutations that fix faster than 99% of neutral fixations** |
| Hard sweeps (#) | Sum of selective sweeps from new mutations |
| Proportion of hard sweeps | Proportion of hard selective sweeps of all selective sweeps |
| Proportion of sweeps from standing | Proportion of selective sweeps from standing variation of all selection sweeps |
| Sweeps (#) | Sum of selective sweeps |
| Sweeps from standing variation (#) | Sum of selective sweeps from mutations that were already segregating before the optimum shift |
| Sweeps/fixations | Ratio of sweeps vs. fixations |
| **Segregating sites** | **Mutations that segregate in the final generation** |
| Max. effect size | Maximal effect size of segregating sites |
| Mean effect size | Mean effect size of segregating sites |
| Mean effect size of negative sites | Mean effect size of segregating sites with negative effects |
| Mean effect size of positive sites | Mean effect size of segregating sites with positive effects |
| Mean frequency of all sites | Mean allele frequency of segregating sites |
| Mean frequency of negative sites | Mean allele frequency of segregating sites with negative effects |
| Mean frequency of positive sites | Mean allele frequency of segregating sites with positive effects |
| Min. effect size | Minimal effect size of segregating sites |
| Negative (#) | Sum of segregating sites with negative effect |
| Positive (#) | Sum of segregating sites with positive effect |
| Proportion of negative sites | Proportion of segregating sites with negative effect of all segregating sites |
| Standard deviation of effect sizes | Standard deviation of effect sizes of all segregating sites |
| Total (#) | Sum segregating sites in the final generation |

